# Wild and cultivated rice host different populations of the blast fungus, *Pyricularia oryzae*, in Mali

**DOI:** 10.1101/2023.10.10.561650

**Authors:** Diariatou Diagne, Henri Adreit, Joëlle Milazzo, Ousmane Koita, Didier Tharreau

## Abstract

Blast is a devastating disease of rice caused by the fungus *Pyricularia oryzae*. The role of infected straw and seed as sources of primary inoculum in blast disease epidemics is well known. The role of alternative hosts is yet to be confirmed. The current study sought to assess if wild rice is a major source of inoculum for cultivated rice by comparing the genetic structure of *P. oryzae* populations from both hosts. Cross infectivity of *P. oryzae* isolates was also assessed using pathogenicity tests. Samples were collected from cultivated and wild rice organs with blast symptoms in irrigated and lowland areas of Mali in Koulikoro, Sikasso, and Ségou regions. Under controlled conditions, *P. oryzae* isolates from wild rice were pathogenic to cultivated rice but, on average, had a narrower range of cultivar compatibility. Results of pathogenicity tests suggest that *P. oryzae* isolates from wild rice have the potential to attack cultivated rice in the field. However, populations of *P. oryzae* on cultivated and wild rice were genetically differentiated. Thus, although physically close, each host plant hosts a different population of the pathogen, and our results support the hypothesis that wild rice is not a major source of blast inoculum for cultivated rice.

## INTRODUCTION

Blast is recognized as one of the most devastating diseases of rice in the world (Savary et al., 2000). The disease is caused by the fungus *Pyricularia oryzae* (synonym *Magnaporthe oryzae*). This model pathosystem for research on host-pathogen interactions (Dean et al., 2012) is also a major challenge for food security (Pennisi, 2010). *Pyricularia oryzae* is an ascomycete fungus that reproduces mainly by asexual multiplication. This widely described asexual cycle is the only mode of reproduction observed in nature. The cycle starts with the adhesion of a spore to the leaf, followed very quickly (in less than two hours) by the formation of a germ tube. The appressorium formed at the tip of the germ tube allows penetration into the plant cell. One to two weeks after infection, *P. oryzae* is able to sporulate and start a new cycle (Ribot et al., 2008). Rice blast is distributed on all continents and in every area where rice is grown (Ou, 1985). Globally, despite the use of control methods, losses are estimated at 4% of the annual world rice harvest (Savary et al., 2019) and represent a lack of staple food for millions of persons.

The genetic structure of *P. oryzae* populations infecting rice has been extensively studied on a global scale to understand the evolution of the pathogen and to adapt control techniques, in particular, the deployment of varietal resistance management strategies. Characterization of *P. oryzae* populations has been carried out at different geographic scales with different molecular markers and genome sequences (Adreit et al., 2007; Gladieux et al., 2018a; Tharreau et al., 2009; Thierry et al., 2022). Studies encompassing samples from different continents are consistent with clustering in four main genetic groups, each with a wide large geographical distribution, and with Asian isolates represented in all four groups (Gladieux et al., 2018a; Thierry et al., 2022). Although, traces of ancestral recombination were identified (Gladieux et al., 2018a, Thierry et al., 2022), populations of the blast fungus on rice are clonal, with the potential exception of populations that could reproduce sexually in very limited areas of Asia (Saleh et al., 2012; Thierry et al., 2022). These studies also showed that Asia is the center of diversity and the origin of the pathogen populations (Thierry et al., 2022). However, little data exist on the diversity and population structure of the pathogen in Africa. Mutiga et al. (2017) studied the genetic diversity of *P. oryzae* rice populations and evaluated the virulence spectrum of this pathogen in West Africa and East Africa. These authors identified seven genetic groups and showed differentiation between West and East African populations. Kassankogno et al. (2017) showed that populations from the two neighboring countries, Burkina Faso and Togo, are differentiated but that some multi-locus genotypes are shared, suggesting some migration between these countries. A recent study (Odjo et al., 2021) confirmed this hypothesis and showed the presence in Africa and Madagascar of the four genetic groups previously described in Asia (Gladieux et al., 2018a; Thierry et al., 2022). Populations in West Africa, East Africa, and Madagascar were highly differentiated (Odjo et al., 2021).

Like any disease, knowing inoculum sources is necessary to develop sustainable control methods. Evidence that wild species are sources of inoculum for cultivated plants are scarce and often indirect for aerial fungal pathogens. The disease severity of ergot on brome grasses and in neighboring fields of barley suggests that brome grasses could be a source of inoculum for barley (Wyka and Broders, 2022). On the contrary, several studies show the divergence of fungal pathogen populations on different hosts. Modern populations of *Rhynchosporium communis* originate from but are differentiated from a population on a wild grass (Brunner et al., 2007). Stukenbrock et al. (2011) showed that the divergence of the wheat-adapted pathogen *M. graminicola* from an ancestral population infecting wild grasses in the Middle East occurred about 10,500 years ago. *Pyrenophora teres f. teres* (Ptt) show divergence of populations on different host populations. Populations from different hosts have distinct evolutionary histories and do not share genotypes. In addition, Ptt isolates could not infect different hosts, confirming host specificity (Linde and Smith, 2019). Linde et al. (2016) assessed and compared the diversity of *Rhynchosporium commune* on weedy and cultivated barley using microsatellites. The study showed that barley is an important auxiliary host of *R. commune*, harboring highly virulent pathogen types capable of transmission to barley.

For blast disease, some primary inoculum sources are well known, such as infected seed and infected straw (Guerber and TeBeest, 2006; Hubert et al., 2015; Long et al.; 2001; Raveloson et al., 2013, 2018). On the other hand, the role of weeds as inoculum sources is controversial. Based on pathogenicity test in controlled conditions, some weeds have been presented as sources of inoculum for blast epidemics on rice, including *Digitaria sanguinalis* (Choi et al., 2013), *Festuca arundinacea*, *Lolium multiflorum*, *Anthoxanthum odoratum*, *Phalaris arundinacea* (Kato et al., 2000), *Rottboellia exaltata, Echinochloa colona*, and *Leersia hexandra* (Mackill and Bonman, 1986). But, to date, this hypothesis has not been confirmed by observations in the field. On the contrary, genetic (Borromeo et al., 1993) and phylogenetic studies (Gladieux et al., 2018b) have shown that strains causing epidemics on rice belong to a single clade, which is not sampled on other hosts (with the exception of a barley sample from Thailand). Similarly to weeds, certain species of wild rice were suspected of serving as alternative hosts. In Africa, the role of wild rice as an inoculum source for Asian cultivated rice was demonstrated for two important diseases, Bacterial leaf streak (Wonni et al., 2014) and RYMV (Traoré et al., 2009). Pathogenicity tests of *P. oryzae* isolates from wild rice (*O. meridionalis*) in northern Australia showed that many local rice varieties of *O. sativa* are susceptible (Khemmuk et al., 2016). On the contrary, although on a limited number of samples (10 isolates), *P. oryzae* population from wild rice (*O. rufipogon*) in Cambodia showed a different pathogenicity spectrum than populations from cultivated rice (Fukuta et al., 2014). This latter study suggests that both pathogen populations may be differentiated. So, to date, there is no evidence that wild species of rice are inoculum sources for blast epidemics on cultivated rice.

Isolating *P. oryzae* from wild rice and showing that these isolates are pathogenic on cultivated rice are not sufficient to demonstrate a role as inoculum source. Both hosts could act as separated compartments and epidemics could be independent, i.e., without demographic and genetic mutual impact on blast populations. Such an example of “parallel” epidemics was observed for blast populations on indica and japonica varieties in the Yuanyang terraces of Yunnan Province in China (Liao et al., 2016). *P. oryzae* isolates from indica varieties were genetically different from isolates from japonica varieties, although both types of varieties were cultivated in close proximity and although isolates from indica varieties had the potential to attack the cultivated japonica varieties. Mali offers an opportunity to address this question since *O. sativa*, the Asian rice (cultivated worldwide) cohabits with the African rice *O. glaberrima* (cultivated in some parts of West Africa), as well as with other wild rice species such as *O*. *longistaminata* and *O. barthii*. Wild rice is usually found in drains, dikes and dams that are close to cultivated rice fields. In that context, wild species of rice could be sources of inoculum for cultivated Asian rice. We took advantage of this specific context to test if the *P. oryzae* populations on both hosts were genetically similar. Our working hypothesis was that, if wild rice is a major source of inoculum for cultivated rice, then there should be important gene flows between *P. oryzae* populations on both hosts and these populations should be genetically similar (i.e. poorly differentiated). We also performed pathogenicity tests in controlled conditions to evaluate the infectivity on *O. sativa* of *P. oryzae* isolates collected on *O. longistaminata*.

## MATERIAL AND METHODS

### Sampling and conservation of samples

Sampling was carried out on leaves, panicle nodes (neck), and stems of cultivated and wild rice showing blast symptoms. They were collected from irrigated and lowland areas between 2017 and 2019, at different stages of rice development in three Regions (Koulikoro, Sikasso, and Ségou) in Mali. In total, 9 sites were surveyed (Sélingué, Manikoura, Niéna, Baguineda, M’pegnesso, Niono Nango Sahel, Niono N3-Bis, Niono N7, Loulouni Faraka Banakoro; Fig. 1; Supplementary Table 1). Diseased organs were collected on different plants distributed in the plots, avoiding to take several plants in disease focus. The number of diseased organs collected per plot depended on the incidence in the field. Access to wild rice was very difficult to assess disease incidence because wild rice was found in water-filled drains and is rarely seen on dikes. A sample was defined as a set of diseased organs collected from the same plot and on a single variety at a single date. On average, the estimated size of a plot is 1/4 hectare. We isolated several strains per sample but mainly one per plant (a maximum of 3 diseased organs were collected per plant).

**Figure 1:**
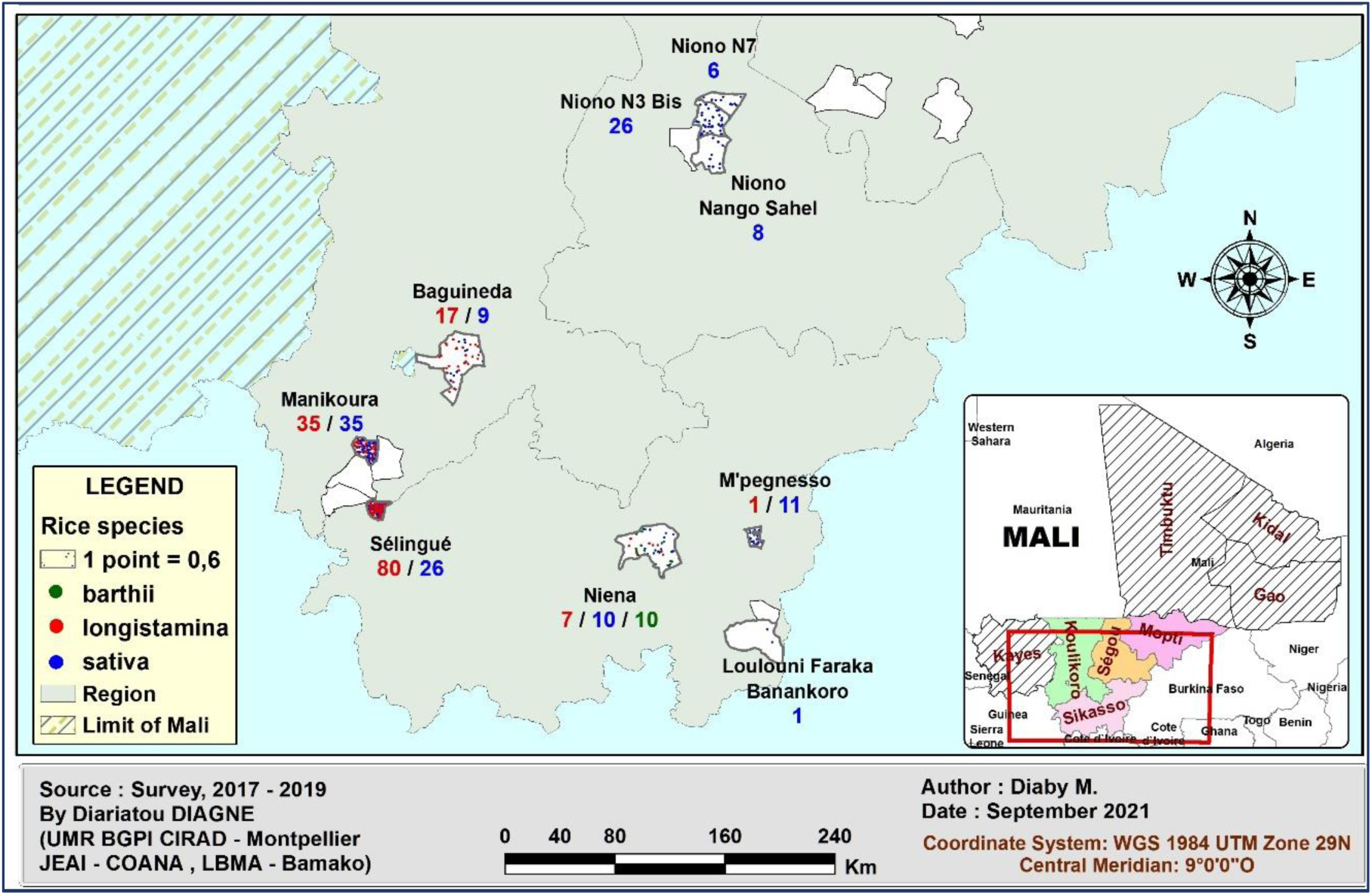
Mapping of *P. oryzae* strains isolated from *O. longistaminata* (red), *O. barthii* (green) and *O. sativa* (blue) by site and species from 2017 to 2019.

### Isolation and conservation of isolates of *P. oryzae*

Diseased organs were placed on a moistened filter paper disc in sterile Petri dishes and incubated at 25°C for 24 h. The fragments were then observed under a binocular microscope (Olympus SZ-PT) to assess the presence or absence of *Pyricularia* conidia. The conidia were collected with a glass needle and deposited on the surface of bacto-agar medium (45 g of bacto-agar in 1 L water). The Petri dish was sealed with tape (Tesa® masking tape 50 m x 19 mm) and incubated at 25°C for 24 h for germination. A germinated conidium was placed on rice flour medium (agar 15 g, rice flour 20 g, yeast extract 2.5 g, water 1 L, and 500,000 units of Penicillin G added after autoclaving) and incubated at 25°C. A single strain was isolated per plant sampled. For storage, an actively growing mycelium plug was deposited on a sterilized filter paper (Whatman N°5) placed on rice flour agar medium (Silué and Nottéghem, 1990). Five to seven days later, the mycelium-covered filter paper was removed and placed in a sterile empty Petri dish with a lid and incubated at 35°C for four days for drying. It was then cut into small fragments and placed in sterile paper bags, placed in a plastic sheath, and sealed under vacuum for conservation at - 20°C (Valent et al., 1986).

A total of 282 strains from different varieties were isolated between 2017 and 2019 from 9 locations (Table 1; Supplementary Table 2). Eleven Malian strains collected between 1986 and 2011 from *O. sativa* were included in the study as controls. These strains were previously characterized by Odjo et al., (2021).

**Table 1:**
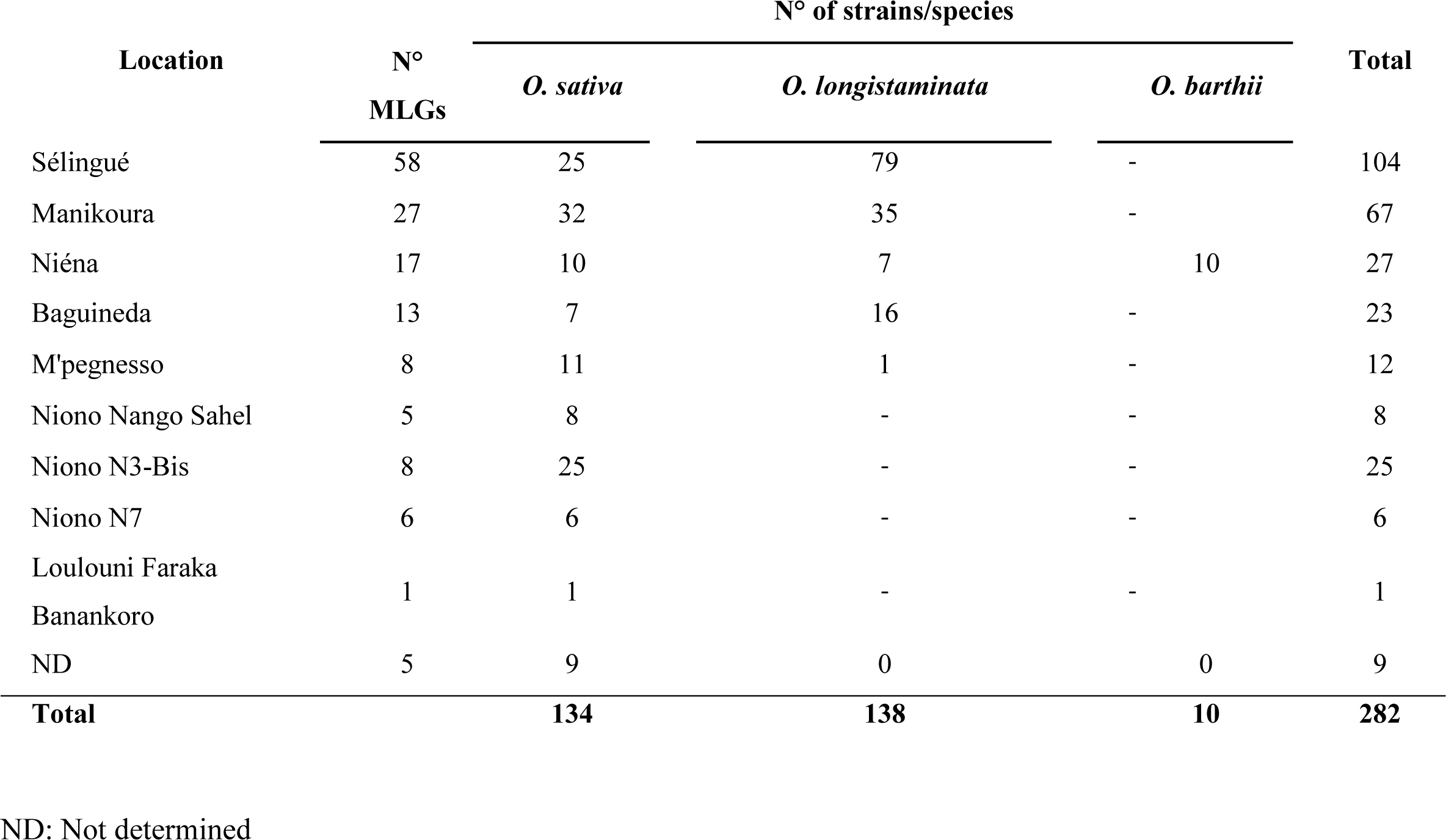
Distribution of isolates by sites and host species and number of multi-locus genotypes per site.

### DNA extraction and microsatellite amplification

After four days of culture in rice flour liquid medium (rice flour 20 g, yeast extract 2 g, water 1 L), the mycelium was recovered in Eppendorf tubes, and genomic DNA was extracted according to the protocol described in Gladieux et al., (2018a).

A total of 282 isolates (271 from the 2017-2019 sampling and the 11 reference strains) were genotyped with 12 Simple Sequence Repeats (SSR) markers (Supplementary Tables 3 & 5) previously used for *P. oryzae* genotyping (Adreit et al.; 2007; Odjo et al.; 2021; Saleh et al.; 2012). These markers were amplified by PCR (QIAGEN Multiplex PCR kit) as previously described (Saleh et al., 2012) with a total volume of 5 μL, including 2.5 µL of Master Mix, 0.5 µL of 10x Mix primers, 0.5 µL of 5x Q solution, and 1.5 µL of genomic DNA (10 ng/µL). The PCR program was as follows: i) predenaturation at 95°C for 15 min, ii) denaturation at 94°C for 30 min, iii) annealing at 57 to 63°C for 90 min, iv) extension at 72°C for 60 min, v) repetition of steps 2 to 4 for 40 cycles, and vi) final extension at 72°C for 30 min. The resulting products were separated and analyzed on a 16-capillary ABI Prism 3130XL machine (Applied Biosystems, Foster City, USA), and the amplicons were evaluated for size by fluorescence measurement. For this analysis, 1.5 μL of amplified products (1/70 dilution) were mixed with 15 μL Formamide HiDi and GeneScan-500LIZ size marker (Applied Biosystems). Amplifications were repeated twice and control strains were included in all amplification plates to verify calibration. When inconsistent results were obtained between two technical replicates, amplification was performed a third time. The isolates with more than two missing data were excluded from the analysis.

### Analysis of genotyping data

The raw data collected were analyzed and transcribed into allele sizes using the GeneMapper V4.1 (Applied Biosystems) tool. We defined an MLG as a unique combination of alleles across all loci genotyped. Strains were grouped in Multi-Locus Genotypes (MLGs) using a script developed by Sébastien Ravel (Odjo et al.; 2021; https://pypi.org/project/MLG-assign/). The first strain of the genotyping datasheet was assigned to MLG1. The second strain was assigned to MLG1 if its genotype was identical to the genotype of the first strain at all loci. The second strain was assigned to MLG2 if its genotype was different for at least one allele at one locus. The procedure was repeated until the genotype of each strain was compared to the MLGs previously identified. Genotypes with missing data may be 100% similar to more than one MLG. In this case, we chose not to assign that genotype to any MLG. Thus, not all isolates could be assigned to an MLG. A discriminant analysis of principal components (DAPC) was performed using the "adegenet" package of the R software (v3.2.2) (Jombart, 2008; Jombart and Ahmed 2011; R Core Team, 2017). DAPC analysis was performed to cluster, without a priori, the MLGs into genetic groups. The mean number of alleles per locus (*Na*) and the unbiased genetic diversity (*H_n.b_*_.;_ Nei, 1987) were calculated using Genetix v4.05 (Belkhir et al., 2004). The genetic diversity of *P. oryzae* populations on wild and cultivated rice in Mali was analyzed. The mean number of private alleles (*N_p_*) was estimated as the number of alleles present in a single genetic group, combining all markers. Genetic differentiation between populations was estimated with the *F_ST_* index (Weir and Cockerham 1984) calculated with Genepop version 4.2 (Raymond and Rousset 1995; Rousset, 2008). We analyzed the genetic differentiation between *P. oryza*e populations on wild (*O. longistaminata*) and cultivated (*O. sativa*) rice within each of the four genetic groups.

### Pathogenicity testing of wild rice isolates on cultivated rice

The pathogenicity of 20 Malian isolates of *P. oryzae* collected from 2017 to 2019 was assessed. Among them, 12 isolates were collected from wild rice *O. longistaminata*, 4 isolates from wild rice *O. barthii,* and 4 isolates from cultivated rice *O. sativa*. Isolates were selected to represent the genetic diversity observed (Table 2).

**Table 2:**
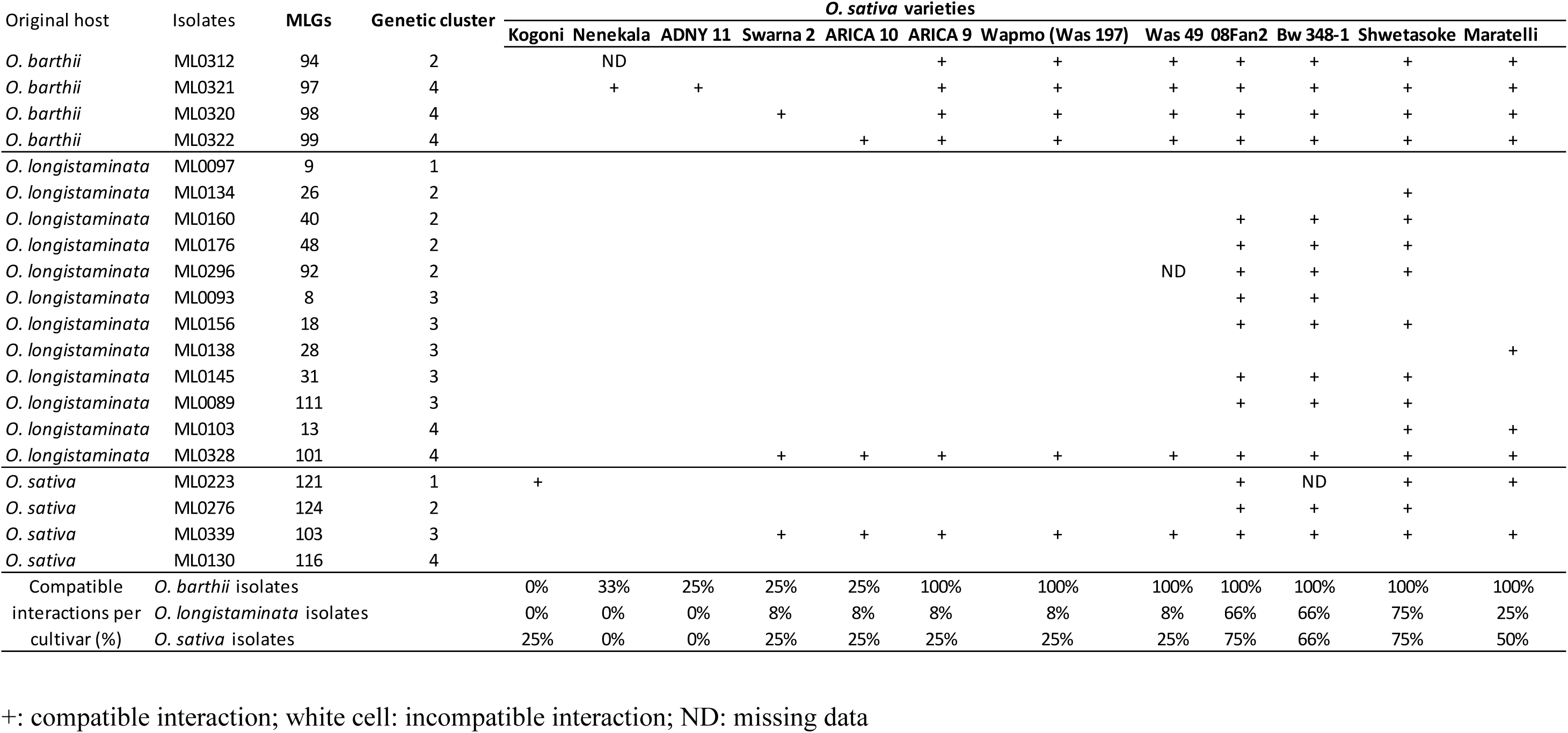
Pathogenicity test on cultivated varieties.

Isolates were inoculated on 11 certified varieties of *O. sativa* cultivated in Mali [08Fan2, ADNY 11, ARICA 10, ARICA 9, Bw 348-1, Kogoni, Nenekala, Shwetasoke, Swarna 2, Wapmo (Was 197), Was 49], and *O. sativa* susceptible check variety Maratelli. The 12 accessions were sown in peat soil (Jiffy compost, reference 284795) in a 33 x 45 x 5 cm plastic tray and cultivated in the greenhouse (temperature set points of 26°C during the day and of 21°C at night; humidity between 50 and 70%). Inoculations were carried out 28 days after sowing (4- to 5-leaf stage) by spraying 30 mL per tray of a spore suspension (Gallet et al., 2016) and were repeated three times. *Pyricularia oryzae* isolates were cultured for one week on rice flour agar medium following the conditions described above for isolate conservation. Spores were collected by flooding the plate with 5 mL of water and scraping the medium. The spore suspension was adjusted to 25,000 spores/mL and 0.5% gelatin. The inoculated plants were incubated at 25°C and 100% relative humidity for one day and then transferred for seven days to a growth chamber (12 h photoperiod; 26°C during the day, 21°C at night). Disease symptoms were assessed seven days after inoculation. The scoring was done according to the 1 to 6 scale described by Gallet et al. (2016). Scores 1 to 3 were considered as incompatible reactions (resistant variety), whereas scores 4 to 6 were considered as compatible reactions (susceptible variety). When data from the three replicates were inconsistent, the corresponding strain x accession interaction was considered as missing data (Table 2).

## RESULTS

*O. longistaminata* was commonly found in the localities of Sélingué (Sikasso region), Manikoura, and Baguineda (Koulikoro region), whereas *O. barthii* was found only in Niéna (Sikasso region). Blast was observed and sampled in all these sites and on all species present. There was no blast disease on *O. longistaminata* and *O. barthii* in Niono (Fig. 1). To compare *P. oryzae* populations on wild and cultivated rice in Mali, we carried out s analyses at two geographic scales (whole country/localities) and at two genetic levels of the pathogen population (genotype/genetic clusters). Because *P. oryzae* populations on rice are clonal in most of the rice-growing areas, including Africa, most population genetic analyses we conducted were based on multilocus genotypes rather than on allelic frequencies.

### Population diversity and structure at the whole country scale

The genetic diversity was similar for *P. oryzae* populations sampled on wild rice (*O. longistaminata*) and on cultivated rice (*O. sativa*) at the whole country scale. The unbiased genetic diversity (*H_nb_*) was 0.62 and 0.58, and the mean number of alleles per locus (*N_a_*) was 9 and 8, respectively (Table 3). The two populations were moderately differentiated, as indicated by a significant but relatively low *F_ST_* value (0.16) for *P. oryzae* (Saleh et al., 2014). The average number of private alleles (*N_p_*) was 3.1 and 2.2, respectively (Table 3). These relatively high average numbers of private alleles (Np) also supported differentiation between the two populations. The sample size (n=10) of the population sampled on *O. barthii* was considered too small to calculate parameters of genetic diversity and differentiation.

**Table 3:** Genetic diversity of *P. oryzae* populations on wild and cultivated rice in Mali.

| Host | N | Diversity parameters |  |  |
| --- | --- | --- | --- | --- |
| | | $H_{nb}$ | $N_a$ | $N_p$ |
| *O. longistaminata* | 138 | 0.62 | 9 | 3.1 |
| *O. sativa* | 134 | 0.58 | 8 | 2.2 |
| **Total** | **272** |  |  |  |

N: total number of individuals, $H_{nb}$ : unbiased gene diversity; $N_a$ : mean number of alleles per locus, $N_p$ : mean number of private alleles per locus.
We used the 12 SSR listed in Supplementary Table 3 for the genotyping.

Two-hundred-fifty-four out of 282 isolates genotyped were unambiguously assigned to 125 multi-locus genotypes (MLGs). Eighty-three MLGs were represented by a single isolate (hereafter named single MLGs), and 42 MLGs were represented by two or more isolates (hereafter named multi MLGs; Table 4). Among these latter, 35 were sampled only on a single host: 18, 14, and 3 on Asian cultivated rice (*O. sativa*) and on wild rice species *O. longistaminata* and *O. barthii*, respectively. There was a limited number of shared MLGs between hosts since only six were sampled on both *O. sativa* and *O. longistaminata* and one on both *O. longistaminata* and *O. barthii* (Table 4).

**Table 4:** Distribution of multi-locus genotypes by species.

| MLGs | Species-specific |  |  | Shared |  |  |
| --- | --- | --- | --- | --- | --- | --- |
|  | *O. sativa* | *O. longistaminata* | *O. barthii* | *O. sativa/O. longistaminata* | *O. sativa/O. barthii* | *O. longistaminata/O. barthii* |
| Single MLGs* | 36 | 47 | 0 | - | - | - |
| Multi- MLGs | 18 | 14 | 3 | 6 | 0 | 1 |

\* Single MLGs: MLGs represented by one isolate; Multi MLGs: MLGs represented by two or more isolates.
5 We used the 12 SSR listed in Supplementary Table 3 for the genotyping.

DAPC analysis was carried out to cluster MLGs in genetic groups without a priori, i.e., using only the genotypic information of the strains. To determine the number of clusters (K), variation of the BIC value was calculated for varying values of K. BIC value regularly decreased with increasing K values (Supplementary Fig. 1). Because the curve did not show a marked inflection, we could not determine an optimal K value based on this approach. Instead, we applied the recommendation of the DAPC tutorial described by Jombart and Collins (2015) and tried to find the value of K that “better summarizes the data than others”. The graphic display of DAPC (Fig. 2) results showed that the genotypes were clearly separated into a minimum number of three groups (Fig. 2). Additional analysis of DAPC results (Fig. 2; Supplementary Fig. 2 & 3) showed that clustering in more than five groups seemed to create an unnecessary (overlapping groups) and unreliable (assignation of genotypes changing from one run to the other) level of clustering. Hence four genetic clusters (named G1, G2, G3, and G4) were defined.

**Figure 2.**
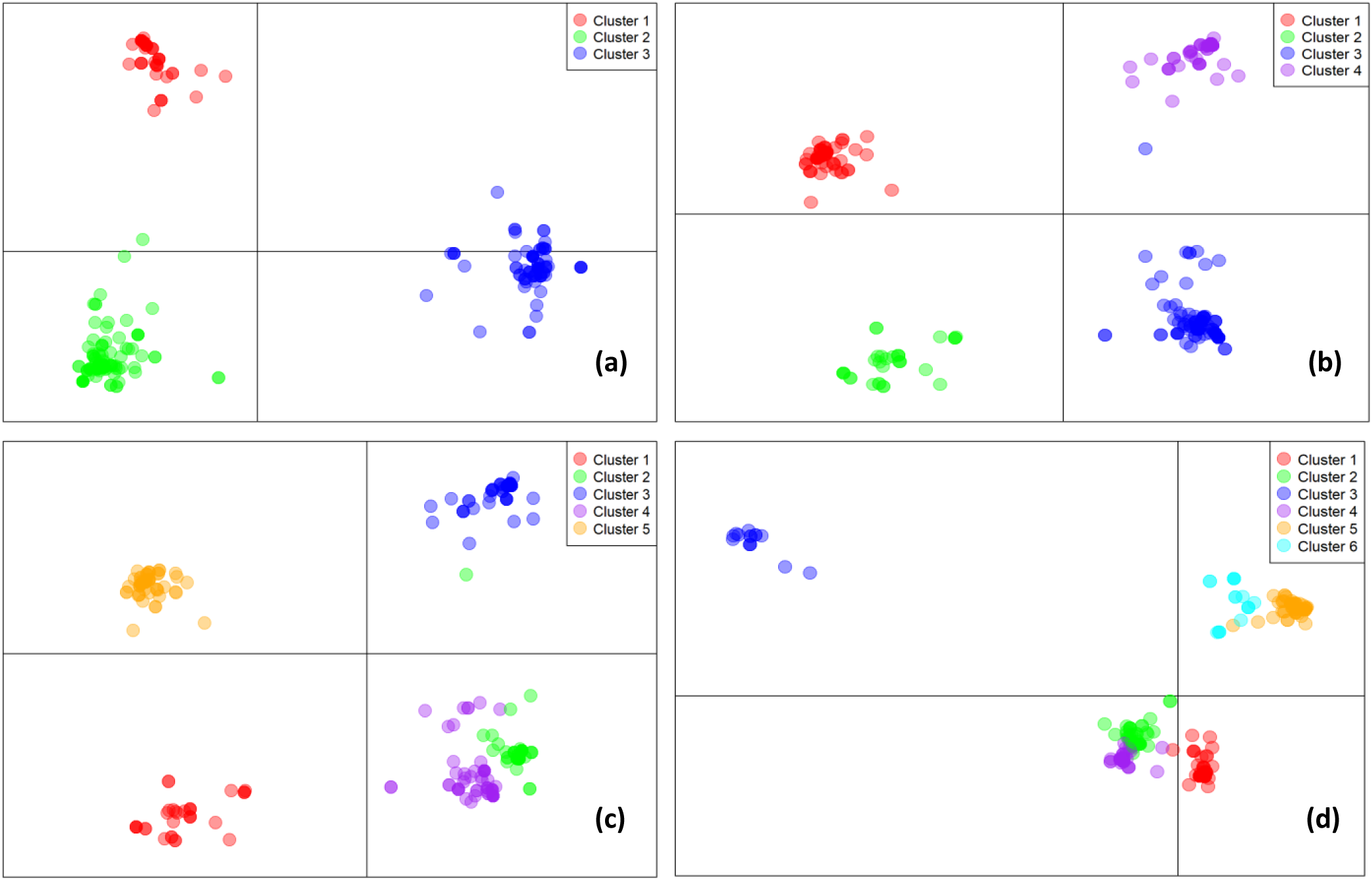
Clustering of genotypes by the DAPC method with the number of cluster (K) varying between three (a), four (b), five (c) and six (d). Each color represents a different cluster.

As expected, and because genotypes were clustered based on genetic similarity, *F_ST_* values were high between the different genetic groups (Table 5). We then analyzed the genetic differentiation between *P. oryzae* populations on wild *O. longistaminata* and on cultivated *O. sativa* rice within each of the four genetic clusters. We observed that the populations isolated from *O. longistaminata* and *O. sativa* were significantly differentiated. *F_ST_* indices between the two populations ranged between relatively low to average values (0.41, 0.16 and 0.31 for G1, G2, and G3, respectively, Table 5). On the contrary, for the G4 cluster, populations from *O. longistaminata* and *O. sativa* were poorly differentiated (*F_ST_* = 0.02; Table 5). A bias due to the small sample size could explain this exception.

**Table 5:** *F_ST_* statistics between genetic clusters of *P. oryzae* populations on wild and cultivated rice in Mali.

| Host | Host Genetic cluster | *O. longistaminata* |  |  |  | *O. sativa* |  |  |
| --- | --- | --- | --- | --- | --- | --- | --- | --- |
|  |  | 1 | 2 | 3 | 4 | 1 | 2 | 3 |
| *O. longistaminata* | 2 | 0.65 | - | - | - | - | - | - |
|  | 3 | 0.64 | 0.66 | - | - | - | - | - |
|  | 4 | 0.46 | 0.59 | 0.51 | - | - | - | - |
| *O. sativa* | 1 | 0.41 | 0.72 | 0.73 | 0.66 |  | - | - |
|  | 2 | 0.64 | 0.16 | 0.63 | 0.47 | 0.76 | - | - |
|  | 3 | 0.65 | 0.65 | 0.31 | 0.50 | 0.73 | 0.62 | - |
|  | 4 | 0.44 | 0.54 | 0.49 | 0.02 | 0.71 | 0.43 | 0.49 |

5 Values for populations from the same genetic cluster but from different hosts are highlighted 6 in grey.

### Population diversity and structure at the local scale

To compare populations at the local scale, we focused on Sélingué, Manikoura, Niéna, and Baguineda localities, where the sample size was sufficient for comparisons between populations isolated from wild rice and cultivated rice (Fig. 1; Table 1).

In Sélingué, 10 single MLGs and 2 multi MLGs (representing a total of 9 isolates) were specific of cultivated rice *O. sativa* (Figure 3a). Similarly, we identified 30 single MLGs and 11 multi MLGs (30 isolates) specific to wild rice *O. longistaminata* (Figure 3a, Supplementary Table 4). Four MLGs were shared between both species. All four Genetic Clusters (GCs) were identified on *O. sativa* and *O. longistaminata.* Clusters G1 was dominant on *O. sativa* (12/25 isolates), whereas G2 (37/79 isolates) and G3 (27/79 isolates) were dominant on *O. longistaminata* (Table 6).

**Figure 3:**
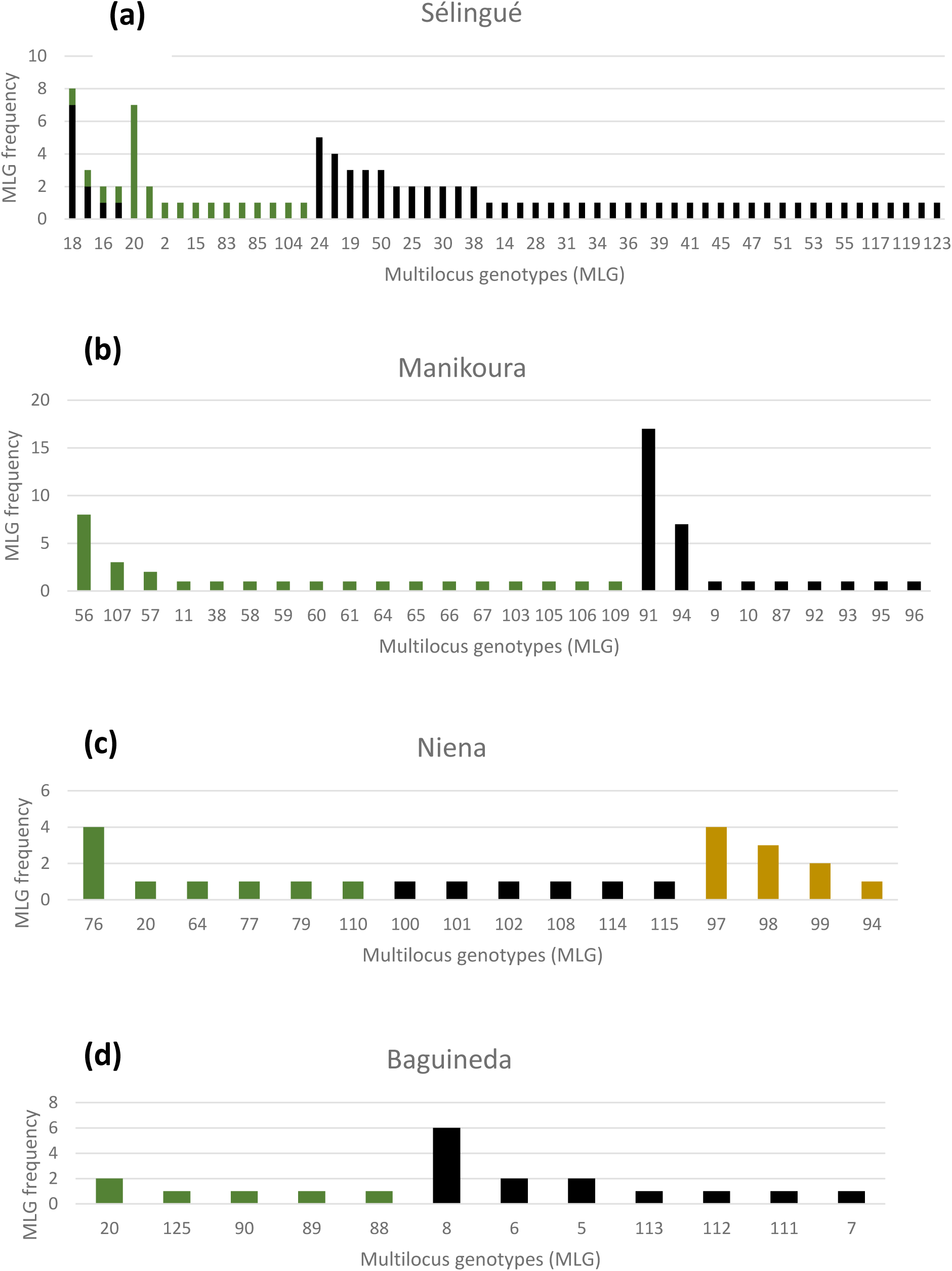
Distribution of MLGs by site and species. Number of isolates per multi-locus genotype in four localities: (a) Sélingué, (b) Manikoura, (c) Niena, and (d) Baguineda. Black: *O. longistaminata*; Green: *O. sativa*; Mustard: *O. barthii*.

**Table 6:**
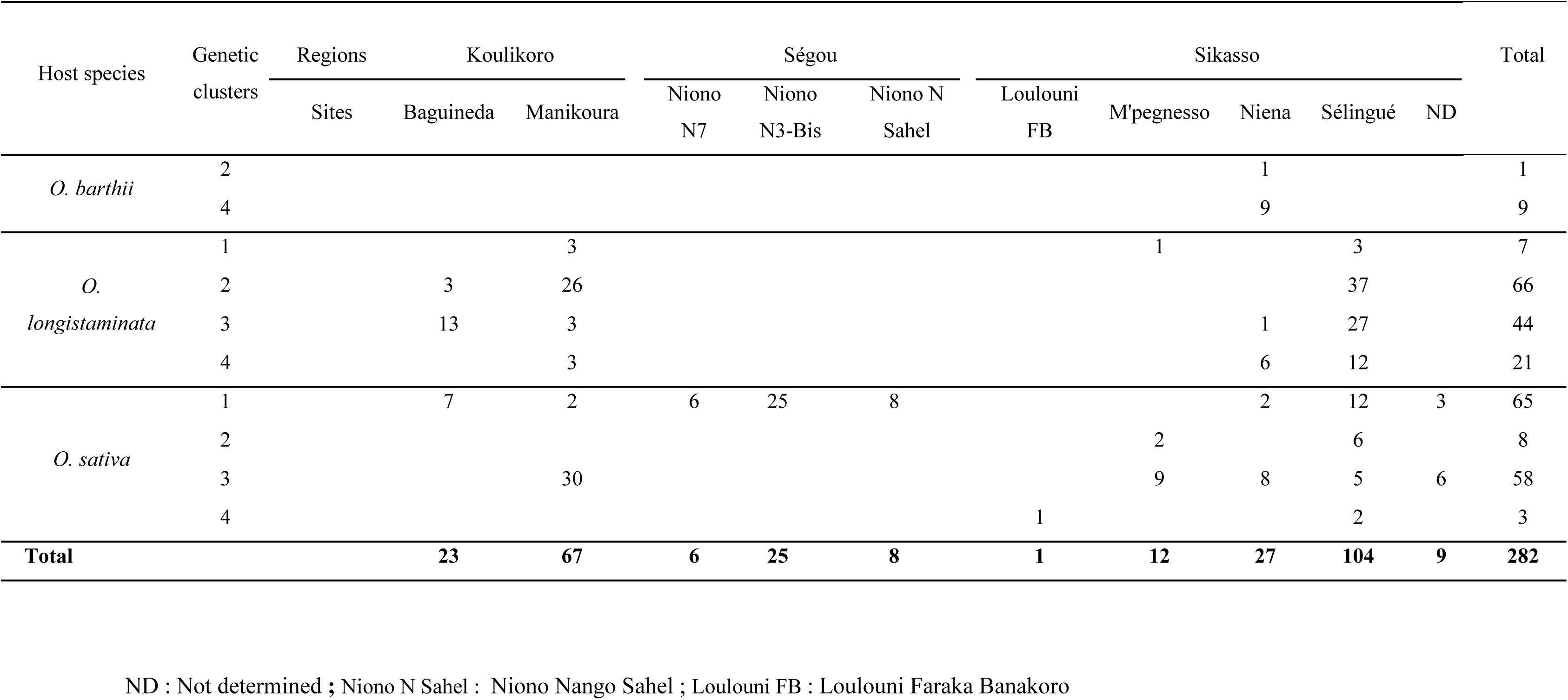
Distribution of genetic clusters by species and sites. Number of *P. oryzae* isolates per site, host species and genetic clusters.

In Manikoura, 14 single and 3 multi MLGs (13 isolates) were specific to *O. sativa*, and 7 single and 2 multi MLGs (24 isolates) were specific to *O. longistaminata* (Figure 3b, Supplementary Table 4). There was no genotype shared between wild and cultivated rice in Manikoura. Genetic clusters G1 and G3 were sampled on *O. sativa*, G3 being more represented (30/32 isolates), whereas the four genetic clusters were identified on *O. longistaminata* with G2 as the dominant one (26/35 isolates; Table 6).

In Niéna, 5 single and 1 multi MLGs (4 isolates) were specific of *O. sativa*, 1 single and 3 multi MLGs (9 isolates) were specific to the wild rice *O. barthii*, and 6 single MLGs were specific of *O. longistaminata* (Figure 3c, Supplementary Table 4). There was no genotype shared between wild and cultivated rice. A dominance of cluster G4 on the two wild species in Niéna was observed. G2 (1/10 isolates) and G4 (9/10 isolates) were found on *O. barthii,* while G3 (1/7 isolates) and G4 (6/7 isolates) were found on *O. longistaminata* (Table 6). G1 (2/10 isolates) and G3 (8/10 isolates) were sampled on *O. sativa*.

In Baguineda, we identified 4 single and 1 multi MLGs (2 isolates) specific of *O. sativa* and 4 single and 3 multi MLGs (10 isolates) specific to *O. longistaminata* (Figure 3d, Supplementary Table 4). There was no genotype shared between wild and cultivated rice in Baguineda. G1 (7 isolates) was the only genetic cluster sampled on *O. sativa*, whereas G2 (3/16 isolates) and G3 (13/16 isolates) were identified on *O. longistaminata* (Table 6).

### Pathogenicity of isolates from *O. longistaminata* on cultivated rice

Most of the varieties (including the susceptible check Maratelli) were incompatible with the majority of isolates, except three varieties (namely 08Fan2, Bw 348-1, and Shwetasoke; Table 2). The average ratio of compatible interactions between *P. oryzae* isolates and *O. sativa* varieties was similar for isolates from *O. longistaminata* and *O. sativa*: 0.24 (range: 0 to 0.75) and 0.34 (range: 0 to 0.75), respectively (Table 2). Isolates from *O. barthii* appeared to have higher frequencies of compatible interactions than isolates from *O. sativa*: 0.69 (range: 0.66 to 0.75). had a. This broad spectrum may be explained by a bias due to the small number of isolates from *O. barthii* tested and probably a narrower genetic diversity because they came from the same place and have closely related genotypes.

## DISCUSSION

The present genetic study shows that *P. oryzae* populations on cultivated and wild rice in Mali are significantly but barely differentiated. This result suggests that, at the country scale, the gene pool shared between both populations is broad. However, comparisons of the clones and genetic clusters composing pathogen populations from each host show a clear differentiation. Each population is mainly composed of specific MLGs. Few MLGs were sampled on both hosts, and an even smaller number were sampled in the same area on both hosts. Although the same four genetic groups are sampled on both hosts, their frequencies are different between hosts. These results show that wild rice (*O. longistaminata*) and cultivated rice (*O. sativa*) in Mali are hosting different populations of the blast fungus. These results do not support the hypothesis that wild rice could be a main source of inoculum for cultivated rice.

Maratelli is considered as a susceptible check because of its broad range of compatibility with *P. oryzae* isolates from rice. Results from different published studies with a very diverse collection of *P. oryzae* isolates from *O. sativa* showed that 94% of the 172 isolates tested were compatible (Gallet et al., 2016; Thierry et al., 2022). In this study, Maratelli was incompatible with an unexpectedly high ratio of isolates (50%). Most of these incompatible isolates (9/10) were isolated on *O. longistaminata*. Malian isolates from wild rice may carry avirulence factors that impede them from infecting Maratelli. Two avirulence genes to Maratelli were already identified and mapped (Mandel et al., 1997). Alleles of these genes triggering an incompatible reaction in Maratelli originated from non-rice *P. oryzae* isolates.

Pathogenicity tests on cultivated rice also revealed that isolates from wild rice are pathogenic to cultivated rice under controlled conditions. In addition, isolates from wild rice did not appear to be less pathogenic on *O. sativa* varieties cultivated in Mali than *O. sativa* isolates, i.e., under controlled conditions, isolates of wild rice (*O. longistaminata*) are pathogenic to a small number of varieties. So, the isolates from *O. longistaminata* have the potential to infect *O. sativa* varieties cultivated in Mali. Similar results were observed with *P. oryzae* isolates from wild rice (*O. meridionalis*) in Australia. These isolates were pathogenic to local rice varieties after artificial inoculation (Khemmuk et al., 2016). Despite this potential, and based on population genetic structure, *P. oryzae* isolates generating epidemics on *O. longistaminata* do not seem to cause epidemics on *O. sativa* in the field. Two non-exclusive causes can be provided to explain this observation. First, our evaluation of compatibility did not take into consideration other pathogenicity components, which may be important in field epidemics (e.g., production of spores by lesion area unit). Second, isolates from *O. longistaminata* are less fit on *O. sativa* than isolates from *O. sativa* and are thus excluded when competing in the field. In *P. oryzae*, fitness differences were observed between populations from two rice subspecies and were hypothesized to contribute to the pathogen population structure (Liao et al., 2016). Additional experiments measuring quantitative differences in pathogenicity and fitness would be needed to test these hypotheses for *P. oryzae* populations of wild and cultivated rice.

## ACKNOWLEDGEMENTS

We thank Youssouf Diarra and the team that collected the samples, specially Mamadou Dembelé of IER Niono at the Office du Niger, the JEAI CoANA team in LBMA, the teams of the Baguineda, Manikoura, Sélingué, Niena, M’pegnesso and Loulouni stations for their contribution. This work was supported by grants of the RICE CRP, a grant from the French Embassy in Mali to Diariatou Diagne, IRD support to Ousmane Koita group (JEAI CoANA), and the support of Institutes of the coauthors (CIRAD, USTT-B). Genotyping data used in this work were produced through the GenSeq technical facilities of the “Institut des Sciences de l’Evolution de Montpellier” with the support of LabEx CeMEB, an ANR "Investissements d’avenir" program (ANR-10-LABX-04-01).

## DATA AVAILABILITY STATEMENT

The data that support the findings of this study are available from the corresponding author upon reasonable request.

## SUPPORTING INFORMATION LEGENDS

Supplementary Figure 1: Variation of bayesian information criterium value with the number of genetic clusters.

Supplementary Figure 2: MLG haplotype network. MLGs clustered in genetic groups 1, 2, 3 and 4 are colored in orange, light blue, blue, light orange, respectively. The construction of the MLG haplotype network into genetic clusters was developed using PHYLOViZ online software. PHYLOViZ is a platform independent JAVA software that allows the analysis of sequence-based typing methods that generate allelic profiles and their associated epidemiological data. https://online.phyloviz.net/index

Supplementary Figure 3: Structure and discriminant analysis of principal components (DAPC) plots of genetic clusters of *Pyricularia oryzae*. (a) Barplot DAPC K=2; (b) Barplot DAPC K=3; (c) Barplot DAPC K=4; (d) Barplot DAPC K=5; (e) Barplot DAPC K=6

Supplementary Table 1: Samples collected per sites from 2017 to 2019. Niono N Sahel: Niono Nango Sahel; Loulouni FB: Loulouni Faraka Banakoro

Supplementary Table 2: Distribution of *P. oryzae* isolates by site and host. Niono N Sahel: Niono Nango Sahel; Loulouni FB: Loulouni Faraka Banakoro

Supplementary Table 3. Primers and polymorphism of the 12 Simple Sequence Repeat (SSR) markers used for genotyping (Odjo et al., 2021) a Number and size of alleles in the African population. b PIC: Polymorphic Information content.

Supplementary Table 3: Distribution of MLGs by site and host species. MLGs: Multi-Locus Genotypes; Unique MLG: MLG represented by a single isolate; ND: Not determined; Niono N Sahel: Niono Nango Sahel; Niono N3-Bis: Niono N3-Bis; Loulouni FB : Loulouni Faraka Banakoro

Supplementary Table 5: Concentrations of SSR markers used for the genotyping.

## Notes

### Competing Interest Statement

The authors have declared no competing interest.

