## Supplementary tables and figures for "Wild and cultivated rice host different populations of the blast fungus, *Pyricularia oryzae*, in Mali"

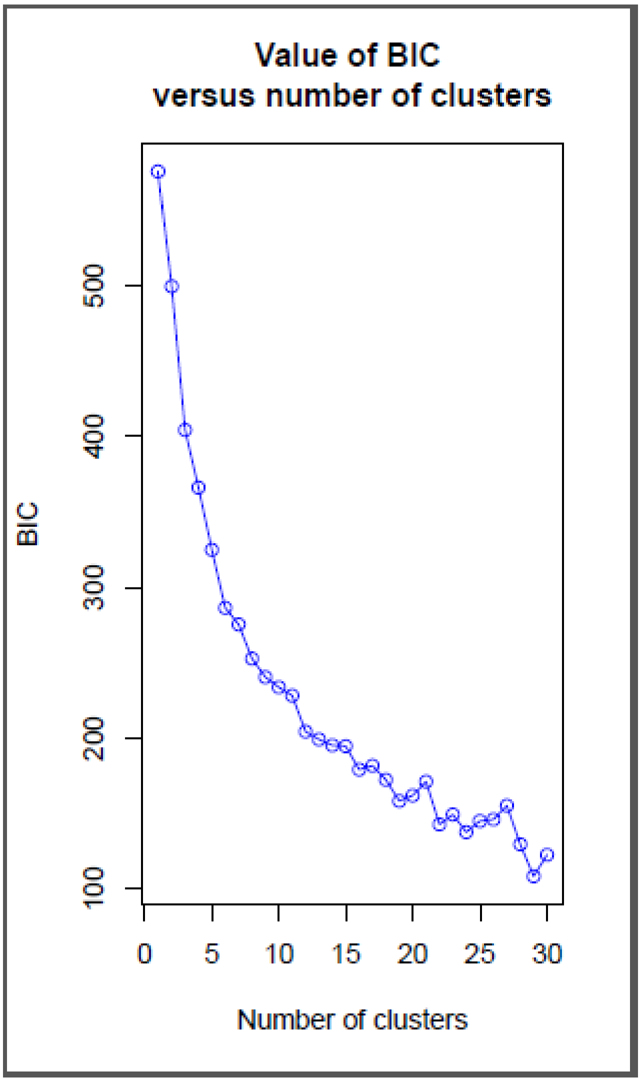


Supplementary Figure 1: Variation of bayesian information criterium value with the number of genetic clusters.

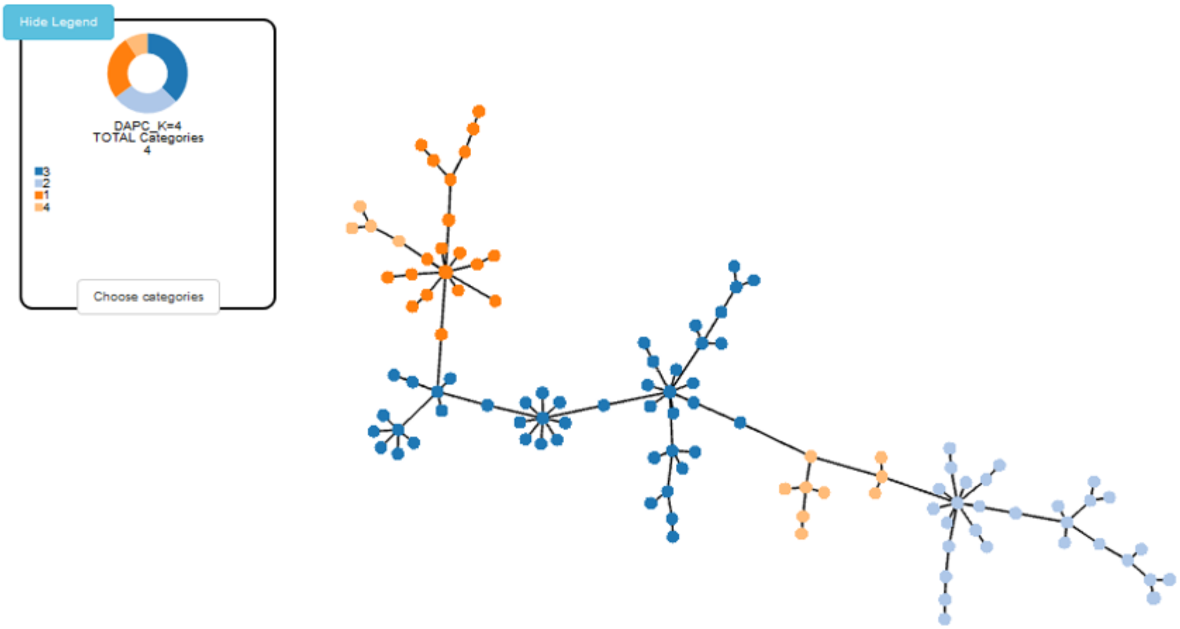


Supplementary Figure 2: MLG haplotype network. MLGs clustered in genetic groups 1, 2, 3 and 4 are colored in orange, light blue, blue, light orange, respectively.

The construction of the MLG haplotype network into genetic clusters was developed using PHYLOViZ online software. PHYLOViZ is a platform independent JAVA software that allows the analysis of sequence-based typing methods that generate allelic profiles and their associated epidemiological data.

https://online.phyloviz.net/index

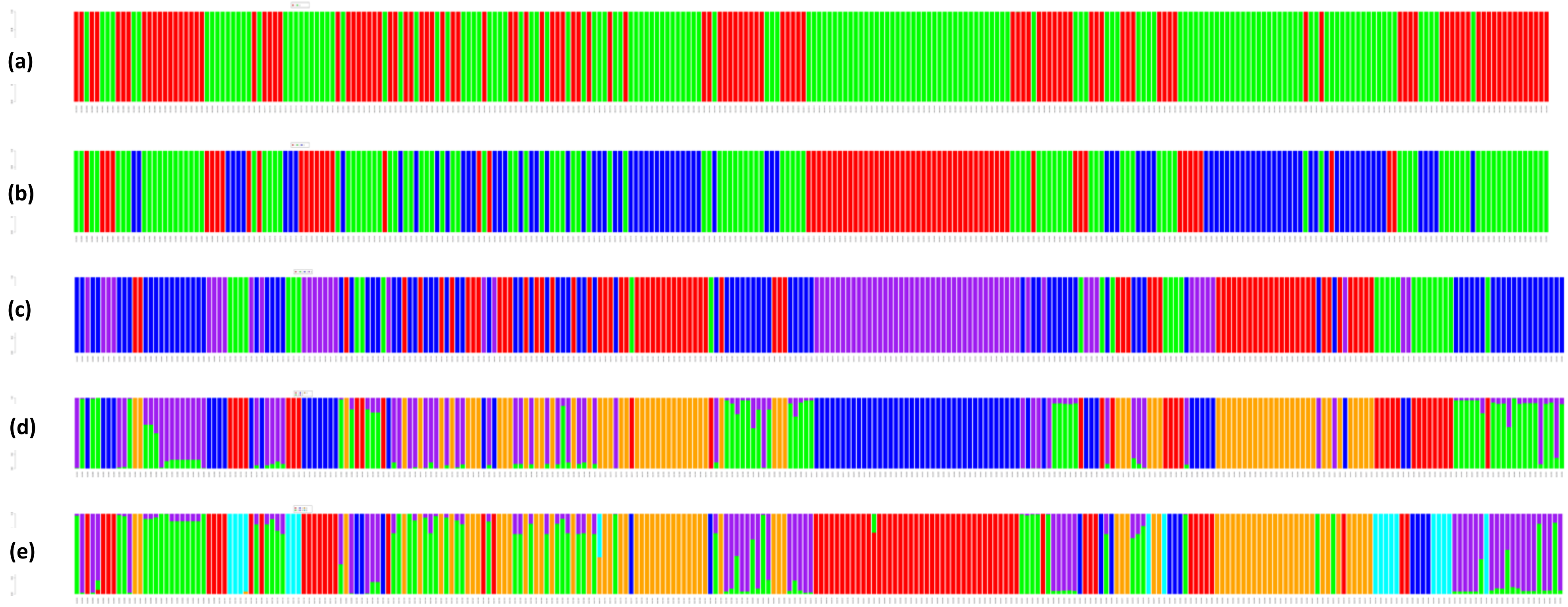


Supplementary Figure 3: Structure and discriminant analysis of principal components (DAPC) plots of genetic clusters of *Pyricularia oryzae*

(a) Barplot DAPC K=2; (b) Barplot DAPC K=3; (c) Barplot DAPC K=4; (d) Barplot DAPC K=5; (e) Barplot DAPC K=6

Supplementary Table 1: Samples collected per sites from 2017 to 2019

| Region | Locality | Collection dates | Total number of plots | Total number of samples | Number of samples processed | Total number of strains collected |
|---|---|---|---|---|---|---|
| Koulikoro | Baguineda | 2017 | 6 | 11 | 11 | 17 |
| | | 2018 | 8 | 14 | 12 | 0 |
| | | 2019 | 3 | 4 | 1 | 9 |
| | Manikoura | 2017 | 13 | 15 | 15 | 4 |
| | | 2018 | 7 | 13 | 4 | 20 |
| | | 2019 | 2 | 4 | 4 | 46 |
| Sikasso | Selingué | 2017 | 14 | 17 | 17 | 22 |
| | | 2018 | 13 | 25 | 23 | 82 |
| | | 2019 | 4 | 4 | 1 | 2 |
| | Niena | 2018 | 7 | 7 | 6 | 10 |
| | | 2019 | 5 | 7 | 3 | 17 |
| | M'pegnesso | 2018 | 3 | 5 | 5 | 12 |
| | Loulouni FB | 2018 | 4 | 6 | 3 | 1 |
| Ségou | Niono N sahel | 2018 | 2 | 3 | 2 | 8 |
| | Niono N3-bis | 2018 | 1 | 2 | 1 | 26 |
| | Niono N7 | 2018 | 3 | 3 | 2 | 6 |
| **Total** | | | **95** | **140** | **110** | **282** |

Niono N Sahel: Niono Nango Sahel; Loulouni FB: Loulouni Faraka Banakoro

Supplementary Table 2**:** Distribution of *P. oryzae* isolates by site and host

| **Varieties/species** | **Region/Location** | | | | | | | | | **Total** |
|---|---|---|---|---|---|---|---|---|---|---|
| | **Koulikoro** | | **Ségou** | | | **Sikasso** | | | | |
| | **Baguineda** | **Manikoura** | **Niono N7** | **Niono N3-Bis** | **Niono N Sahel** | **Loulouni FB** | **M'pegnesso** | **Niena** | **Sélingué** | |
| BW 348-1 (*O. sativa*) | | | | | | | 7 | 2 | | 9 |
| IR32310 (*O. sativa*) | | 33 | | | | | 4 | | 19 | 56 |
| Kogoni 91-1 (*O. sativa*) | 9 | | 6 | 26 | 8 | | | | 7 | 56 |
| Locale (*O. sativa*) | | 2 | | | | | | | | 2 |
| Shwetasoké (*O. sativa*) | | | | | | 1 | | 8 | | 9 |
| *O. barthii* | | | | | | | | 10 | | 10 |
| *O. longistaminata* | 17 | 35 | | | | | 1 | 7 | 80 | 140 |
| **Total** | **26** | **70** | **6** | **26** | **8** | **1** | **12** | **27** | **106** | **282** |

Niono N Sahel : Niono Nango Sahel ; Loulouni FB : Loulouni Faraka Banakoro

Supplementary Table 3. Primers and polymorphism of the 12 Simple Sequence Repeat (SSR) markers used for genotyping (Odjo et al., 2021)

[a] Number and size of alleles in the African population. [b] PIC: Polymorphic Information content.

| Name | Chromosome | Primers | Number of alleles[a] | PIC[b] | Size (pb)[a] | Final concentration |
|---|---|---|---|---|---|---|
| Pyrms63 | 1 | F:TTGGGATCTTCGGTAAGACG<br>R: GCCGACAAGACACTGAATGA | 3 | 0.22 | 149-161 | 0.2 µM |
| Pyrms37 | 4 | F:ACCCTACCCCCACTCATTTC<br>R: AGGATCAGCCAATGCCAAGT | 3 | 0.21 | 197-209 | 0.2 µM |
| Pyrms319 | 2 | F:TAAGACCACTGGCGGAATCT<br>R: GGCTTTGTCTGGTTGTACGG | 3 | 0.24 | 281-287 | 0.2 µM |
| Pyrms657 | 6 | F:ATCAGTCGAACCCACAAAGC<br>R: ATGTGTGGACGAACCAGTCC | 4 | 0.30 | 164-170 | 0.2 µM |
| Pyrms427 | 5 | F:CTGTCACCACAACCAAGACG<br>R: TTGCCCTGATTTGTCAGTCA | 9 | 0.61 | 199-221 | 0.2 µM |
| Pyrms607 | 3 | F:CCCAAGCTCCATAATACGCTAC<br>R:TCCGAGACTCTTTGGATAGCAC | 4 | 0.35 | 272-287 | 0.2 µM |
| Pyrms233 | 5 | F:TGAGATGGACCGCATGATTA<br>R: TTGATGGCAGAGACATGAGC | 6 | 0.33 | 241-268 | 0.2 µM |
| Pyrms47 | 4 | F:TCACATTTGCTTGCTGGAGT<br>R: AGACAGGGTTGACGGCTAAA | 7 | 0.49 | 157-169 | 0.2 µM |
| Pyrms77B | 3 | F:AGGCTCTCTGCCTACGAAGT<br>R: GCTTTCGGCAAGCCTAATC | 7 | 0.44 | 194-226 | 0.2 µM |
| Pyrms83B | 2 | F:GTCTGCCTCGACTCCTTCAC<br>R: GCAAAGTTGTTTGAGCAAGG | 4 | 0.62 | 103-115 | 0.2 µM |
| Pyrms99B | 5 | F:CACCACTTTATGGCGCAGT<br>R: ACCTAGGTAGGTATACATGTTGTT | 11 | 0.53 | 226-313 | 0.2 µM |
| Pyrms409 | 4 | F: TCCCAGTACTTGCCCATCTC<br>R: ATCTCATATCCGTCGGTCGT | 15 | 0.68 | 299-343 | 0.2 µM |

Supplementary Table 3**:** Distribution of MLGs by site and host species

| MLGs | Species | N° of strains | | | | | | | | | | Total |
|---|---|---|---|---|---|---|---|---|---|---|---|---|
| | | Koulikoro | | Ségou | | | Sikasso | | | | | |
| | | Baguineda | Manikoura | Niono N7 | Niono N3-Bis | Niono N Sahel | ND | Loulouni FB | M'pegnesso | Niena | Sélingué | |
| 15 | *longistaminata* | | | | | | | | 1 | | | 1 |
| 15 | *sativa* | | | | | | | | | | 1 | 1 |
| 16 | *longistaminata* | | | | | | | | | | 1 | 1 |
| 16 | *sativa* | | | | | | | | | | 1 | 1 |
| 18 | *longistaminata* | | | | | | | | | | 7 | 7 |
| 18 | *sativa* | | | | | | | | | | 1 | 1 |
| 38 | *longistaminata* | | | | | | | | | | 2 | 2 |
| 38 | *sativa* | | 1 | | | | | | | | | 1 |
| 43 | *longistaminata* | | | | | | | | | | 1 | 1 |
| 43 | *sativa* | | | | | | | | | | 1 | 1 |
| 49 | *longistaminata* | | | | | | | | | | 2 | 2 |
| 49 | *sativa* | | | | | | | | | | 1 | 1 |
| 94 | *barthii* | | | | | | | | | 1 | | 1 |
| 94 | *longistaminata* | | 7 | | | | | | | | | 7 |
| 1 | *sativa* | | | | | | 3 | | | | | 3 |
| 107 | *sativa* | | 3 | | | | | | | | | 3 |
| 11 | *sativa* | | 1 | 1 | 4 | 1 | | | | | 1 | 8 |
| 2 | *sativa* | | | | | | 3 | | | | 1 | 4 |
| 20 | *sativa* | 2 | | 1 | 13 | 3 | | | | 1 | 7 | 27 |
| 21 | *sativa* | | | | | | | | 2 | | | 2 |
| 22 | *sativa* | | | | | | | | 2 | | | 2 |
| 56 | *sativa* | | 8 | | | | | | | | | 8 |
| 57 | *sativa* | | 2 | | | | | | | | | 2 |
| 63 | *sativa* | | | | | | | | 2 | | | 2 |
| 64 | *sativa* | | 1 | | | | | | | 1 | | 2 |
| 68 | *sativa* | | | | | 2 | | | | | | 2 |
| 70 | *sativa* | | | | 1 | 1 | | | | | | 2 |
| 73 | *sativa* | | | | 2 | | | | | | | 2 |
| 76 | *sativa* | | | | | | | | | 4 | | 4 |
| 78 | *sativa* | | | | | | | | 2 | | | 2 |
| 79 | *sativa* | | | | | | | | 1 | 1 | | 2 |
| 82 | *sativa* | | | | | | | | | | 2 | 2 |
| 12 | *longistaminata* | | | | | | | | | | 2 | 2 |
| 19 | *longistaminata* | | | | | | | | | | 3 | 3 |
| 23 | *longistaminata* | | | | | | | | | | 3 | 3 |
| 24 | *longistaminata* | | | | | | | | | | 5 | 5 |
| 25 | *longistaminata* | | | | | | | | | | 2 | 2 |
| 27 | *longistaminata* | | | | | | | | | | 2 | 2 |
| 30 | *longistaminata* | | | | | | | | | | 2 | 2 |
| 33 | *longistaminata* | | | | | | | | | | 2 | 2 |
| 44 | *longistaminata* | | | | | | | | | | 4 | 4 |
| 5 | *longistaminata* | 2 | | | | | | | | | | 2 |
| 50 | *longistaminata* | | | | | | | | | | 3 | 3 |
| 6 | *longistaminata* | 2 | | | | | | | | | | 2 |
| 8 | *longistaminata* | 6 | | | | | | | | | | 6 |
| 91 | *longistaminata* | | 17 | | | | | | | | | 17 |
| 97 | *barthii* | | | | | | | | | 4 | | 4 |
| 98 | *barthii* | | | | | | | | | 3 | | 3 |
| 99 | *barthii* | | | | | | | | | 2 | | 2 |
| Unique MLGs | | 8 | 18 | 3 | 3 | 1 | 2 | 1 | 2 | 8 | 37 | 83 |
| nd | *longistaminata* | 2 | 4 | | | | | | | 1 | 8 | 15 |
| nd | *sativa* | 1 | 5 | 1 | 2 | | 1 | | | 1 | 2 | 13 |
| **Total** | | **23** | **67** | **6** | **25** | **8** | **9** | **1** | **12** | **27** | **104** | **282** |

MLGs: Multi-Locus Genotypes
Unique MLC MLG represented by a single isolate
ND: Not determined
Niono N Sahel : Niono Nango Sahel ; Niono N3-Bis : Niono N3-Bis ; Loulouni FB : Loulouni Faraka Banakoro

Supplementary Table 5: Concentrations of SSR markers used for the genotyping

| **Simplex 1** | Primers | Final Volume uL | **PCR Working Conc uM** | Initial Conc uM | Initial Vol uL | Primer Total Vol Total uL |
|---|---|---|---|---|---|---|
| | Pyrms 47/48 | 500 | **1** | 10 | 50 | |
| | Pyrms 427/428 | 500 | **1** | 10 | 50 | |
| | Pyrms 99B/100 | 500 | **1** | 10 | 50 | |
| | | | | | **Sum** | **150** |
| **Simplex 2** | Pyrms 409/410 | 500 | **1** | 10 | 50 | |
| | Pyrms 657/658 | 500 | **0.5** | 10 | 25 | |
| | Pyrms 77B/78 | 500 | **1** | 10 | 50 | |
| | | | | | **Sum** | **125** |
| **Simplex 3** | Pyrms 63/64 | 500 | **1** | 10 | 50 | |
| | Pyrms 43B/44 | 500 | **1** | 10 | 50 | |
| | Pyrms 83/84 | 500 | **1** | 10 | 50 | |
| | Pyrms 607/608 | 500 | **0.5** | 10 | 25 | |
| | | | | | **Sum** | **175** |
| **Simplex 4** | | | | | | |
| | Pyrms 37/38 | 500 | **0.5** | 10 | 25 | |
| | Pyrms 233/234 | 500 | **0.5** | 10 | 25 | |
| | Pyrms 319/320 | 500 | **0.5** | 10 | 25 | |
| | | | | | **Sum** | **75** |
| **Repass 5** | Pyrms 99B/100 | 500 | **2** | 10 | 100 | |
| | Pyrms 43B/44 | 500 | **2** | 10 | 100 | |
| | | | | | **Sum** | **200** |
| **Repass 6** | Pyrms 63/64 | 500 | **1** | 10 | 50 | |
| | Pyrms 427/428 | 500 | **1** | 10 | 50 | |
| | Pyrms 233/234 | 500 | **1** | 10 | 50 | |
| | Pyrms 37/38 | 500 | **1** | 10 | 50 | |
| | | | | | **Sum** | **200** |
